# A New Variant of Hepatitis A Virus Causing Transient Liver Enzyme Elevations in Mauritius-origin Laboratory-housed Cynomolgus macaques

**DOI:** 10.1101/2022.11.18.515766

**Authors:** Lars Mecklenburg, Rebecca Ducore, Molly Boyle, Andrew Newell, Laura Boone, Joerg Luft, Annette Romeike, Ann-Kathrin Haverkamp, Keith Mansfield, JJ Baczenas, Nick Minor, Shelby L. O’Connor, David H. O’Connor

## Abstract

Hepatitis A virus (HAV) infects humans and non-human primates causing an acute self-limited illness. Three HAV genotypes have been described for humans and three genotypes have been described for non-human primates. We observed transiently elevated liver enzymes in Mauritius-origin laboratory-housed macaques in Germany and were not able to demonstrate HAV by serology and PCR. Using deep sequencing, we have identified a new HAV genotype with 86% nucleotide sequence homology to HAV genotype IV capsid proteins and approximately 80% nucleotide homology to other HAV genotypes. In situ hybridization indicates persistence in the biliary epithelium up to 3 months after liver enzymes were elevated. Vaccination using a commercial vaccine against human HAV prevented reoccurrence of liver enzyme elevations. Since available assays for HAV did not detect this new variant, knowledge of its existence may ameliorate potential significant epidemiological and research implications in laboratories globally.

**Article Summary Line:** A new genotype of hepatitis A virus, that was not identifiable by available diagnostic assays, caused liver enzyme elevations in laboratory-housed Mauritius-origin Cynomolgus macaques.

## INTRODUCTION

Cynomolgus macaques are commonly used for safety assessment of human biopharmaceuticals under development. Such nonclinical investigations typically involve the determination of indicators of liver injury, i.e. alanine aminotransferase (ALT) and glutamate dehydrogenase (GLDH). ALT and GLDH activities, that exceeded the upper limit of the site-specific historical range, were noted in animals from nonclinical studies conducted in Germany and in naïve animals (prior to allocation to study). Increases in ALT and GLDH activities were only observed in animals of Mauritius-origin and not in any animals originating from mainland Asia. These increases in ALT/GLDH activities were transient, normalized within 6 weeks, and only occurred once in any affected animal. Further, increased ALT/GLDH activities were not associated with specific macroscopic or microscopic correlates in the liver or alterations in indicators of hepatobiliary injury and were considered subclinical in that they were not associated with any observations of ill health.

A root cause analysis was conducted with investigation into potential infectious causes including human hepatitis A virus (HAV), hepatitis B virus, hepatitis C virus, hepatitis D virus und lymphocytic choriomeningitis virus. In addition, investigations were conducted for non-infectious causes such as aflatoxins, intoxication by detergents or leachables from toys that were provided for behavioral enrichment. Since none of the investigations generated a positive result, we initiated a massive parallel sequencing approach, using serum samples from animals that were acutely affected by increased liver enzyme activities. This approach yielded a previously unrecognized variant of HAV.

HAV is a non-enveloped RNA virus in the family *Picornaviridae*. The host range for HAV is limited to man and several species of non-human primate (*1*). The human HAV strains appear to have diverged from the simian HAV strains approximately 3,600 years ago (*2,3*). HAV is likely pervasive in many types of wild monkeys, but systematic data about its prevalence is lacking. Wild cynomolgus monkeys in peninsular Malaysia become infected at a rate comparable to that of humans in the same region (*4*). In captive primates, spontaneous infection with HAV has been reported in great apes, Old World monkeys, and New World monkeys.

Cynomolgus macaques that are experimentally infected with HAV demonstrate viral RNA in serum, saliva, and feces at 7 days postinoculation. Biochemical and histological signs of infection are first seen at 15 days postinoculation (*5*), characterized by increased ALT and chronic periportal inflammation (*6*). Infected animals demonstrate seroconversion with the appearance of anti-HAV IgM and IgG antibodies (*6,7*).

This article presents the pattern of increased ALT and GLDH activities in captive cynomolgus macaques, presumably caused by this novel variant of simian HAV, which evaded existing commercially available serologic and PCR detection tests. We describe our root cause investigation and the subsequent discovery of this novel HAV. Finally, information is presented as to how to mitigate viral-induced effects in a laboratory setting.

## METHODS

### Housing, Origin and General Health of Animals

The facility in which this new variant of HAV was discovered, houses cynomolgus monkeys bred in Mauritius and Asia mainland. Animals are socially housed in groups of 2 to 20 with full compliance to the European Convention for the Protection of Vertebrate Animals used for Experimental and Other Scientific Purposes (ETS123) and Directive 2010/63/EU. Before entering the facility, all animals are screened for Simian Retro Virus (serology and PCR), Simian Immunodeficiency Virus (serology), and Simian T-cell Leukemia Virus (serology). Animals from mainland Asia are additionally screened for Macacine herpesvirus 1 (serology). Animals reported in this investigation were either allocated to a nonclinical safety study of a novel pharmaceutical or were colony animals (i.e. animals not yet allocated to an experiment).

### Biomarkers of Hepatocellular and Hepatobiliary Injury

Blood was collected into tubes without anticoagulant. The indicators for hepatocellular injury consisted of alanine aminotransferase (ALT), aspartate aminotransferase (AST), and glutamate dehydrogenase (GLDH) activities and the indicators for hepatobiliary injury consisted of gamma glutamyltransferase (GGT) and alkaline phosphatase (ALP) activities and total bilirubin concentration. These clinical chemistry endpoints were analyzed from serum by the Konelab™ system (Thermo Fisher Scientific Inc, Waltham, MA). The time points for blood collection for these endpoints varied between studies and animals.

All data were entered into an electronic database (Pristima XD, Xybion, Princeton, NJ). Analysis and visualization were conducted using the TIBCO Spotfire platform.

### Histopathology

In three colony animals, a liver biopsy was collected for histopathologic evaluation immediately following detection of increased ALT and GLDH activities. The liver tissue was fixed in 10% neutral-buffered formalin, paraffin blocks were prepared and sectioned, and slices were stained with hematoxylin and eosin. Microscopic examination was performed by a board-certified pathologist on at least two liver samples per animal.

### Deep Sequencing

Blood samples from 6 animals that demonstrated elevated ALT and GLDH activities were used for massive parallel sequencing with a sequence-independent single-primer amplification (SISPA) approach (*8*), that was updated to sequence influenza vRNA from clinical respiratory samples (*9*). A description of the procedure is provided in the **<u>Appendix</u>**.

Sequencing read data were then analyzed with a novel virus discovery pipeline, available at https://github.com/dholab/Mecklenburg-et-al-2022. This pipeline uses scripts from the BBTools suite (http://sourceforge.net/projects/bbmap/) to mask host-specific reads (bbmask.sh), repair read pairing (repair.sh), and trim Illumina adapters (bbduk.sh). The pipeline then removes PhiX, human, and broad metagenomic reads with bbmap.sh, deduplicates reads with dedupe.sh, and merges paired and unpaired reads with bbmerge.sh. Merged, trimmed reads are then assembled *de novo* with SPAdes (DOI: https://doi.org/10.1002/cpbi.102). After removing short and low-complexity contigs from the assembly (bbmask.sh and breformat.sh from BBtools), the pipeline classifies contigs with megablast, classifies unclassified contigs with blastn, and then outputs data for classified and unclassified contigs (DOI 1: https://doi.org/10.1186/1471-2105-10-421; DOI 2: https://doi.org/10.1016/S0022-2836(05)80360-2). These results showed evidence of a novel, Hepatitis A-like virus in all 5 out of 6 animals.

Next, we visually inspected the SPAdes assemblies and generated a consensus sequence for the novel virus with Geneious Prime (Geneious Primer 2021.2.1). To account for excessive depths of coverage across many of the contigs, which may bias a consensus sequence, we used Geneious to reprocess NovaSeq reads by first trimming to remove adapters, low quality sequences, and 19bp from 5’ end of each sequence. Still in Geneious, we then produced synthetic long reads by merging the trimmed reads with bbmerge, and then mapped these reads to NCBI RefSeq # NC_001489, a Hepatitis A virus reference sequence, with the Geneious mapper. To ensure we detect novel virus reads, we required at least 100 bases of overlap between the reads and the reference sequence and tolerated up to 40% base mismatches. We then generated a prototype consensus sequence from the sample with the best coverage, corrected it by mapping reads from that sample onto its consensus using Multiple Sequence Comparison by Log-Expectation align (*10*), transferred annotations from the human HAV NC_001489, trimmed low quality ends, and translated the consensus sequence into amino acids. Finally, we ran protein and nucleotide BLAST on the 7490 bp consensus sequence of the newly identified HAV, which resulted in several matches that were all approximately 20% nucleotide divergent. GenBank accession numbers for matching sequences (*11,12*) were: AB020564.1 (HAV isolate AH1), M14707 (HAV strain HM175), AY644676.1 (HAV isolate CF53/Berne), AY644670.1 (HAV strain SLF88), AB279732.1 (HAV isolate HA-JNG04-90), M34084.1 (Simian HAV capsid protein VP1 strain PA21), AB279735.1 (HAV isolate HAJ85-1), M59286.1 (HAV capsid proteins VP1, VP2, VP3, VP4, P2A strain Cy145), D00924.1 (Simian HAV gene for polyprotein strain AGM-27). A GenBank formatted file of the consensus sequence, its annotations, and its translation is also available at https://github.com/dholab/Mecklenburg-et-al-2022.

### Quantitative reverse transcription PCR for the New Variant of Hepatitis A Virus

A quantitative reverse transcription PCR (qRT-PCR) method was developed to specifically detect the new variant of HAV, referred to as “simian HAV Macaca/Germany/2021”. Briefly, three sets of primers were designed: primer sets for qPCR of simian HAV Macaca/Germany/2021, one endpoint PCR primer pair that should detect human HAV and simian HAV Macaca/Germany/2021 (and presumably some other divergent HAV), and one set of PCR primer pools that can be used for overlapping amplicon genome sequencing. Using the consensus sequence from the sample with the best coverage, we then generated primers in Geneious Prime. We set the primer tool to generate primers for between nucleotide 1000 and 6000 of the consensus to avoid ends, used the human mispriming library that is built into Geneious, and specified an optimal melting temperature of 60°C. To design endpoint PCR primers, we used a multiple sequence alignment method (*10*) to align all sequences from the tree presented in (*13*), and we generated a consensus sequence in Geneious Prime. Finally, to design overlapping PCR primers for sequencing the whole-genome of simian HAV Macaca/Germany/2021, we used PrimalScheme (*14*) with a target amplicon size of 1200 bases. Primer pairs are summarized in the **<u>Appendix</u>**.

For qRT-PCR, we extracted total RNA from serum samples (0.2 ml each) of 18 animals that showed elevated ALT and GDH activities and of 62 animals without liver enzyme elevation. In addition, we extracted total RNA from liver samples taken from 2 animals with elevated ALT and GLDH activities.

### In Situ Hybridization

Formalin-fixed and paraffin-embedded tissue from liver, gall bladder and gastrointestinal tract was collected from 5 animals that had been assigned to the vehicle control group of a toxicity study. Those animals had demonstrated liver enzyme elevations between 4 days and 11 weeks prior to necropsy.

For *in situ* hybridization tissues were sectioned at 4-5 microns on Superfrost plus slides. In situ hybridization to detect simian HAV Macaca/Germany/2021 as well as Mf-PPIB (positive control and tissue quality control) and DapB (negative control) genes was performed using reagents and equipment supplied by Advanced Cell Diagnostics (ACDBio, Hayward, CA) and Ventana Medical Systems (Roche, Tuscon, AZ). The *in situ* hybridization RNAscope® probes where designed based on the simian HAV Macaca/Germany/2021. The hybridization method followed protocols established by ACDBio and Ventana systems using a 3,3’-Diaminobenzidine (DAB) chromogen. Briefly, sections were baked at 60 degrees for 60 minutes and used for hybridization. The deparaffinization and rehydration protocol was performed using a Sakura Tissue-Tek DR5 stainer with the following steps: 3 times xylene for 3 minutes each; 2 times 100% alcohol for 3 minutes; air dried for 5 minutes. Off-line manual pretreatment was conducted in 1X retrieval buffer at 98 to 104 degrees C for 15 minutes. Optimization was performed by first evaluating PPIB and DapB hybridization signal and subsequently using the same conditions for all slides. Following pretreatment the slides were transferred to a Ventana Ultra autostainer to complete the ISH procedure including protease pretreatment; hybridization at 43 degrees C for 2 hours followed by amplification; and detection with 3,3’-Diaminobenzidine chromogen and hematoxylin counter stain.

### Vaccination

After discovery of simian HAV Macaca/Germany/2021, a vaccination program was initiated at the housing facility. For vaccination, a commercial vaccine against inactivated human HAV HM175 strain, genotype IB (Havrix 720 Kinder, GSK) was used. Every newly arriving animals was once vaccinated (0.5 ml per animal administered intramuscular) within 14 days of arrival.

## RESULTS

### Increased ALT and GLDH Activities

Increased ALT and GLDH activities, which exceeded the upper limit of the test site historical range, were identified in serum collected from animals in the pre-dose phase or from vehicle-treated control animals on nonclinical safety studies conducted between October 2020 and October 2021. ALT and GLDH activities in the affected animals were increased up to 11-fold **(Figure 1)**. Importantly, increased ALT and GLDH activities were only noted at one time-point for affected animals despite longitudinal monitoring. Increased ALT and GLDH activities did not correlate with clinical observations or macroscopic or microscopic findings in the liver at terminal necropsy; were infrequently associated with concurrently increased AST activity (only noted in 6% of the affected animals); and were unaccompanied by increases in indicators of hepatobiliary injury. Further, no similarly increased ALT and GLDH activities were noted in cynomolgus monkeys of Asia mainland-origin in studies concurrently conducted at the facility.

**Figure 1:**
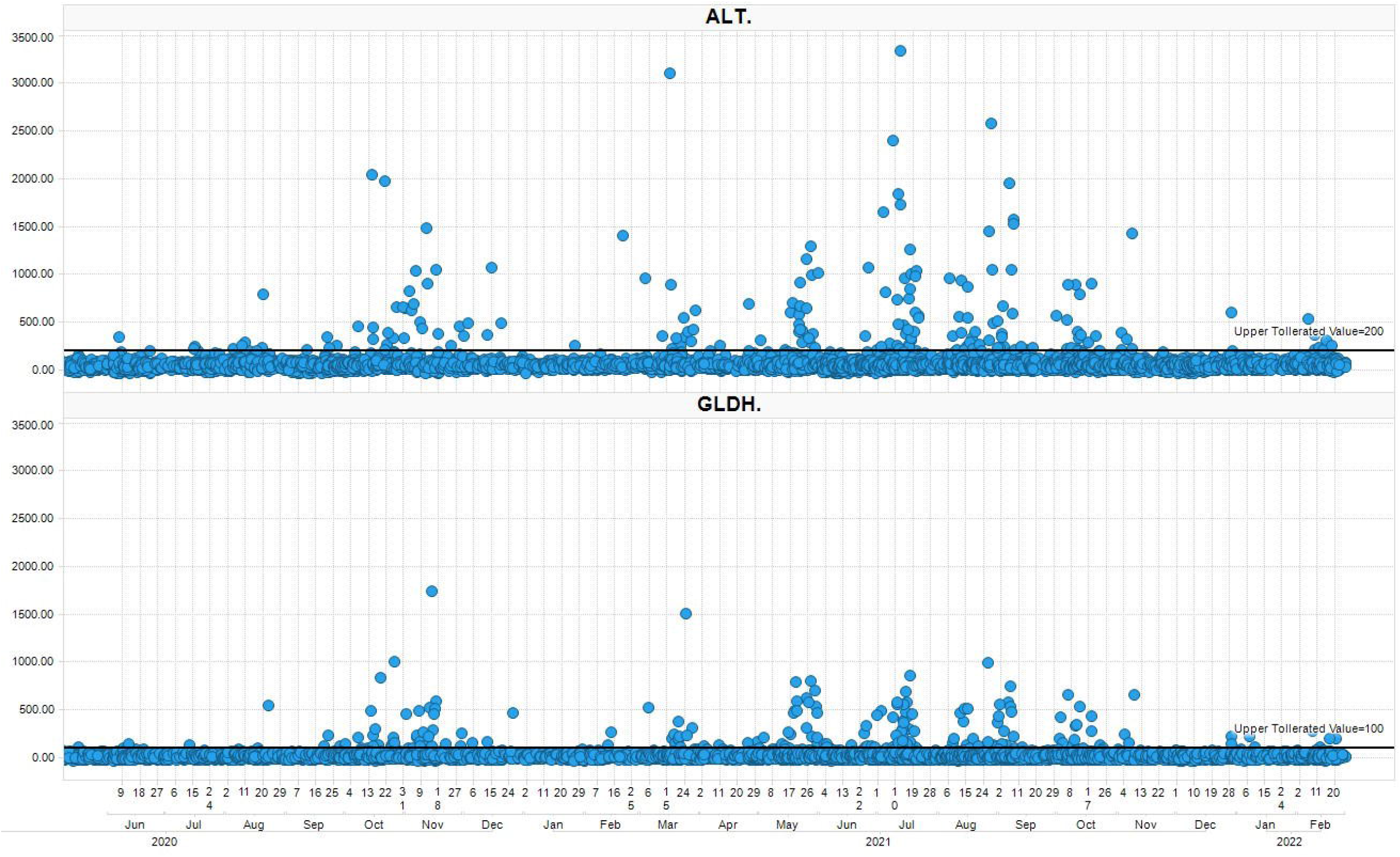
Scatter plot of ALT and GLDH values over time in blood samples from Mauritius origin cynomolgus monkeys assigned to nonclinical safety studies. June 2020 to February 2022. Animals were administered various test articles and vehicles. Upper reference range shown by black line (200 U/l for ALT and 100 U/l for GLDH). Each dot represents a measurement recorded by calendar date.

Screening of liver enzymes from Mauritius-origin animals that had not yet assigned to a study was initiated. Out of 3,912 blood samples collected between December 2020 and March 2022, 144 samples (3.6%) demonstrated increased ALT and GLDH activities **(Figure 2A)**. These were identified between March and October 2021. Sequential monitoring of the ALT and GLDH activities in affected animals indicated that the increased enzyme activities were transient and generally returned to values within the historical range within 6 weeks **(Figure 2B)**.

**Figure 2.**
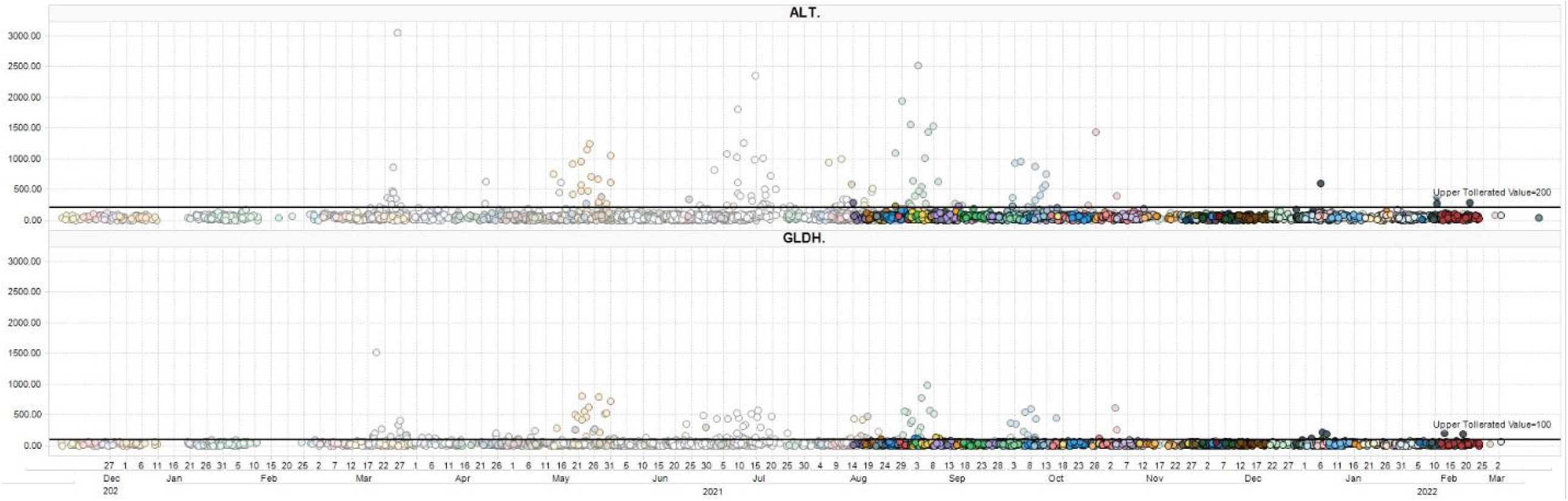
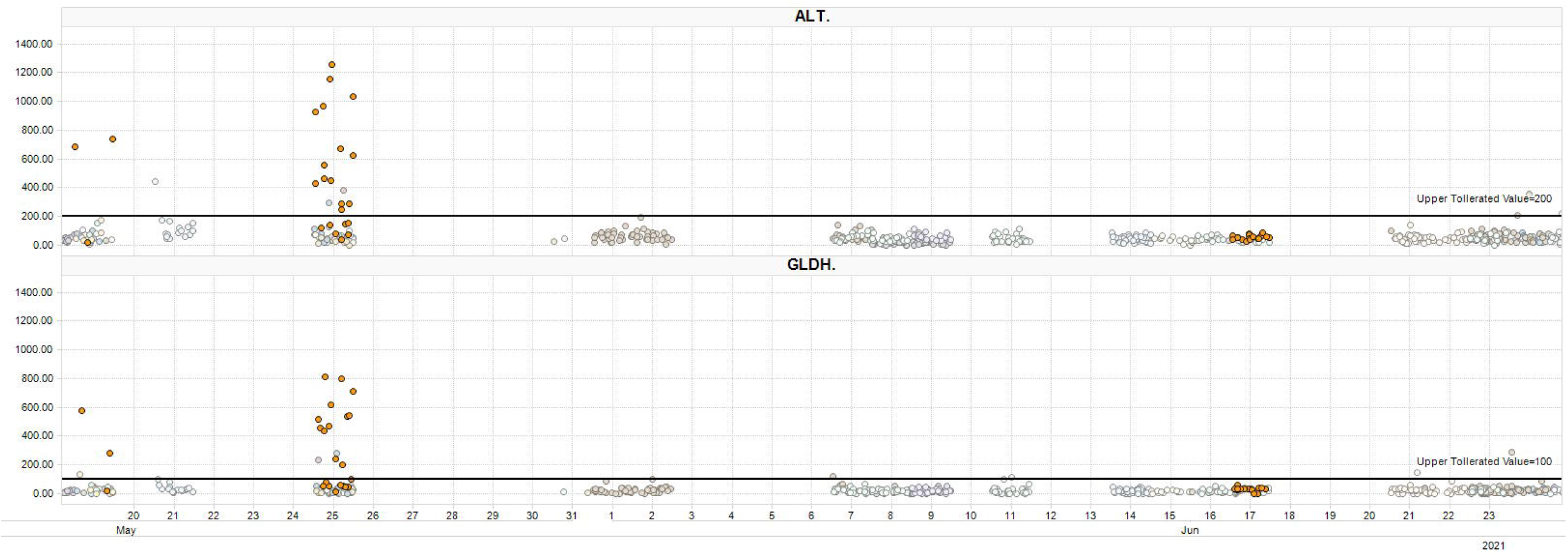
Scatter plot of ALT and GLDH values over time in blood samples taken from naïve Mauritius-origin cynomolgus monkeys. Each dot represents a blood sample; each color represents a cohort of animals that was received at the test site. **(A)** December 2020 to March 2022. Faded colors represent animals that were not vaccinated against Hepatitis A virus, full colors represent animals that were vaccinated upon arrival. **(B)** May 2021 to June 2021. Orange dots represent one specific cohort of animals that was followed up over time.

### Histopathology

In order to verify any histomorphological evidence of chronic periportal inflammation as described (*6*), liver tissue was sampled from three colony animals shortly after detection of increased ALT and GLDH activities. In all animals, microscopy was characterized by minimal to slight periportal mononuclear cell infitrates **(<u>Appendix</u>)**. Such infiltrates of minimal up to slight magnitude are occasionally seen in macaques (*15*). In addition, very few degenerating hepatocytes or hepatocellular single cell necrosis was seen in two animals **(<u>Appendix</u>)** and minimal pigment in macrophages or Kupffer cells (Perl’s positive; Fouchet negative) and minimal hepatocellular vacuolation was seen in one case.

### Deep Sequencing

Massive parallel sequencing yielded in 5 out of 6 samples sequencing reads that match a Hepatitis A-like virus, referred to as simian HAV Macaca/Germany/2021. Consensus sequences of the viruses were virtually identical between samples. The identified nucleotide sequence of simian HAV Macaca/Germany/2021 consists of 7490 bp with 86.08% homology to the simian HAV capsid proteins sequence (*16*) (GenBank accession no. M59286.1). Simian HAV Macaca/Germany/2021 shows 81.41% nucleotide sequence homology to human HAV subtype IIIB genomic RNA (GenBank accession no. AB279735.1), and less than 81% sequence homology to human HAV genotypes IA, IB, IIA, IIB, and IIIA and simian HAV genotype V (GenBank accession no. D00924.1) **(Table 1)**.

**Table:** Nucleotide identity (%) between simian HAV Macaca/Germany/2021 and known HAV genotypes.

At predicted amino acid level, the sequence of simian HAV Macaca/Germany/2021 has 92.00% homology with HAV polyprotein derived from olive baboons (GenBank accession no. ANJ65975.1), 91.97% homology with simian HAV polyprotein derived from a captive rhesus monkey (GenBank accession no. ABX55994.1), and approximately 90% homology with numerous HAV polyprotein sequences from other HAV genotypes obtained from human patients. A comparison of the predicted amino acid sequence specifically for viral protein 1 (nucleotides 2203-3099) revealed a 98.66% identity with partial HAV polyprotein from HAV genotype IV (16) (GenBank accession no. AAA45473.1) **(Figure 3)**.

**Figure 3:**
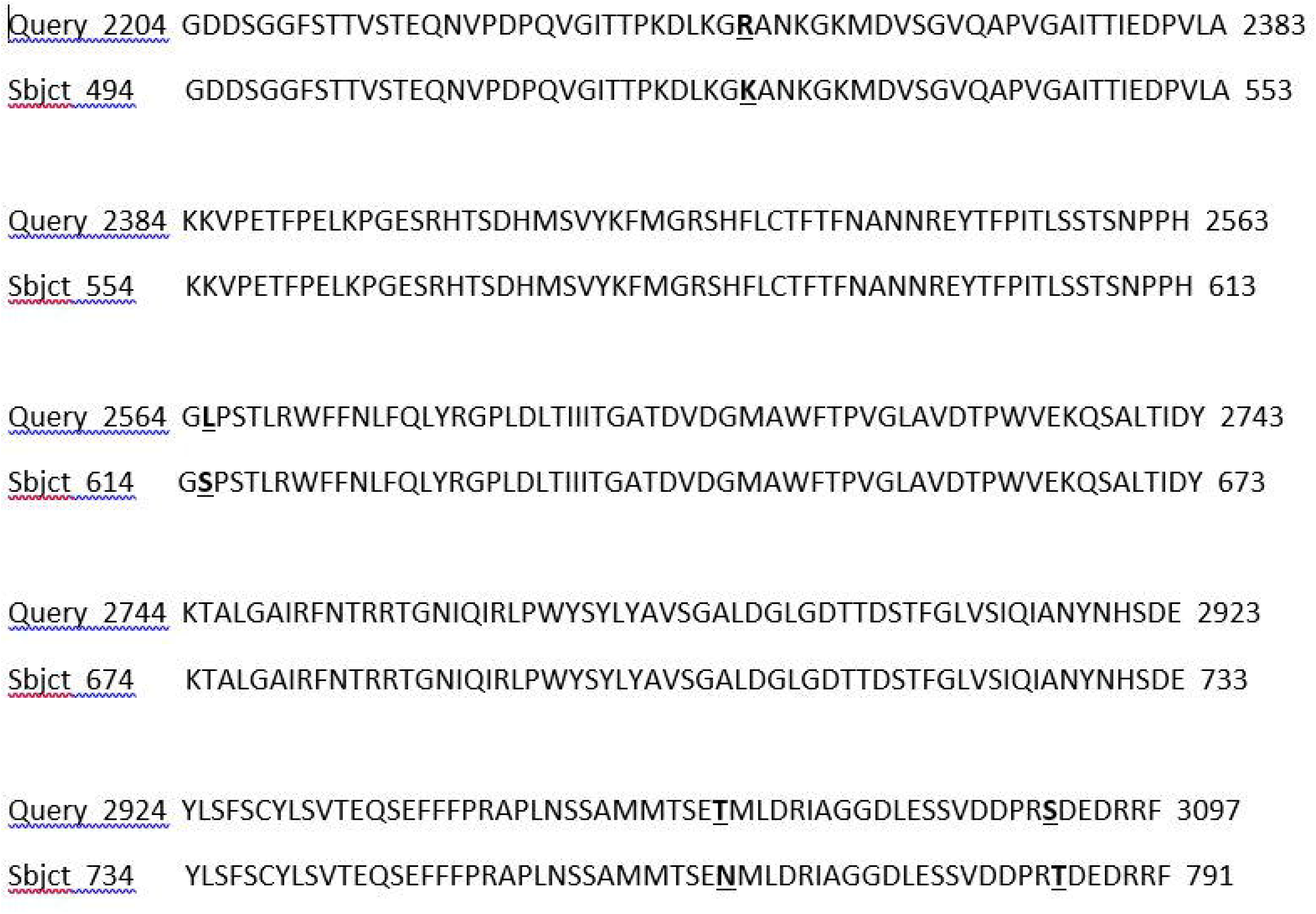
Amino acid sequence alignment between predicted simian HAV Macaca/Germany/2021 viral protein 1 (Query) and HAV polyprotein GenBank accession no. AAA45473.1, range 494 to 791 (Sbjct). Differences are highlighted in bold and underlined.

### Diagnostic PCR

Since conventional serology and PCR assays for human HAV had not detected the new HAV variant, we developed a quantitative RT-PCR method specific for simian HAV Macaca/Germany/2021. This method was used on serum samples from 80 animals and on liver tissue samples from 2 animals. The method detected simian HAV Macaca/Germany/2021 in all 18 serum samples and in both liver tissue samples from animals with increased ALT and GLDH activities. The 62 serum samples from animals without increased ALT and GLDH activities were negative.

### In Situ Hybridization

In situ hybridization detected simian HAV Macaca/Germany/2021 in liver tissue from 5 out of 5 animals. Signal was present to varying degrees within hepatocytes, sinusoidal cells and biliary epithelium **(Figure 4)**. In addition, simian HAV Macaca/Germany/2021 signal was observed in the GALT of one animal with high hepatic viral load. In three animals euthanized 4 days after increases in ALT and GLDH were detected, signal was observed in hepatocytes, sinusoidal lining cells including Kupffer cells and biliary epithelial cells in the liver. In two animals euthanized 11 weeks after increases in ALT and GDLH activities were detected, signal was largely absent from the liver and not present in hepatocytes. Despite an absence of simian HAV Macaca/Germany/2021 signal in hepatocytes of these animals, signal was observed in gall bladder epithelium suggesting the biliary epithelium may represent a reservoir of infection and shedding after resolution of hepatocellular infection.

**Figure 4:**
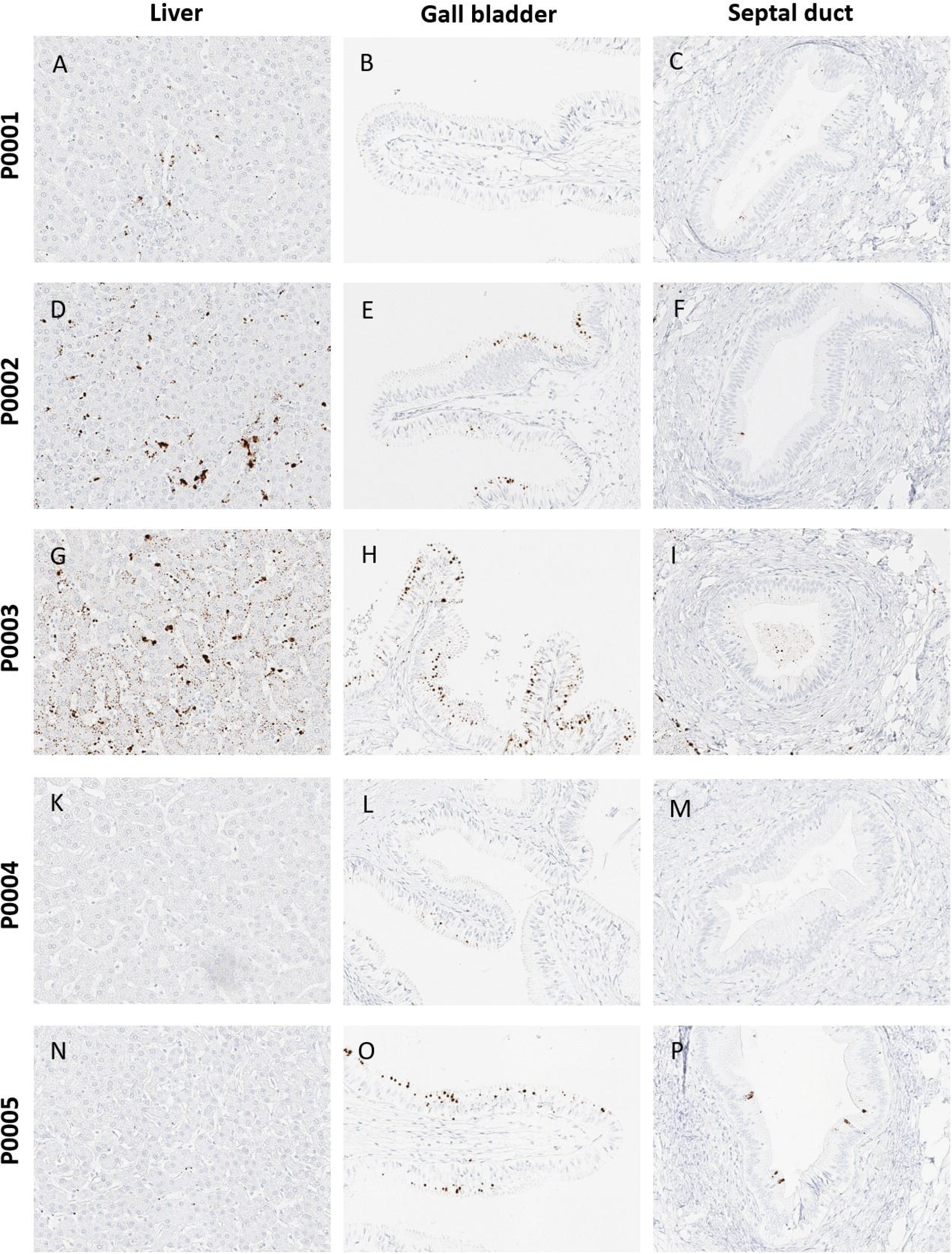
In situ hybridization for MueHAV in tissue from liver (A, D, G,K and N), gall bladder (B, E, H, L and O) and hepatic septal ducts (C, F, I, M and P) in animals euthanized 4 days (P0001, P0002 and P0003) and 11 weeks (P0004 and P0005) after detection of elevations in ALT and GLDH.

### Vaccination

After identification of simian HAV Macaca/Germany/2021, we initiated a vaccination program that uses a commercial vaccine against human HAV GBM strain. All animals newly arriving to the test facility (both of Mauritius and Asia mainland origin) were vaccinated within 14 days of arrival. In addition, all colony animals housed at the facility but not yet allocated to an experiment were also vaccinated. The vaccination program started in August 2021. Since its initiation, newly arriving animals have been monitored with respect to liver enzyme activities. In all vaccinated animals, ALT and GLDH activities have remained within the reference range **(Figure 2A)**.

## DISCUSSION

HAV is a known cause of hepatitis and elevated liver enzyme activities in captive cynomolgus monkeys (*6*). However, disease outbreaks in captive monkeys are rarely reported. At the facility where this new variant of HAV was discovered, no such disease outbreak had occurred for more than 40 years. Moreover, all employees were routinely vaccinated against human HAV, since the virus represents a potential zoonotic risk.

When the increased ALT and GLDH activities were identified, HAV was the main differential diagnosis. Appropriate diagnostic tests (PCR and serology) were initiated but ultimately failed to detect simian HAV Macaca/Germany/2021. Despite broad-ranging diagnostic investigations, no etiology could be identified. Therefore, a massive parallel sequencing approach, also known as next-generation sequencing, was employed. This approach allows identification and characterization of bacteria, fungi, parasites, protozoa, and viruses without prior knowledge of a specific pathogen.

The approach revealed a previously unreported Hepatitis A virus with less than 87% whole genome nucleotide sequence homology to any previously reported HAV. Highest nucleotide sequence homology was detected for HAV capsid proteins VP1, VP2, VP3, VP4, and P2A (*16*). This sequence was derived from two Cynomolgus macaques in the United States: One animal was imported from the Philippines and had serological as well as histopathological evidence of hepatitis, the other was imported from Indonesia and showed clinical signs of a HAV infection.

HAV is a small non-enveloped ancient virus with a long evolutionary history that is distinct from other picornaviruses. The HAV genome is a single-stranded, positive-sense RNA molecule, approximately 7500 nucleotides in length, that serves directly as messenger RNA for translation of HAV-encoded proteins (*17*). Like other picornaviruses, HAV shows a high mutation rate, resulting in the emergence of a genetically diverse cloud of mutants (*18*).

The HAV genome contains an open reading frame encoding a large polypeptide with multiple cleavage sites. It is generally agreed that the polyprotein is processed to four structural capsid proteins, i.e. viral protein (VP) 1 to 4, and seven non-structural proteins (2ABC, 3ABCD). Processing is performed by the proteinase encoded in the 3C region. The host humoral response against the capsid protein represents the primary weapon to control the infection. Capsid proteins VP1 and VP3 are major antibody-binding sites (*18*). While there is good evidence that most human strains of HAV are closely related antigenically, simian strains have significant antigenic differences from human HAV strains (*19*). The newly discovered simian HAV Macaca/Germany/2021 is sufficiently different from known human and simian HAVs, such that diagnostic methods that are directed against human HAV would not detect this new simian variant. This was confirmed by directly comparing the genomic sequence of simian HAV Macaca/Germany/2021 against the PCR primer sequences from the commercially available PCR test used to detect human HAV.

Despite the difference between simian HAV Macaca/Germany/2021 and human HAV, we successfully vaccinated animals with a commercial vaccine against inactivated human HAV genotype IB. This vaccination program was able to prevent further occurrence of increased ALT and GLDH activities.

It is interesting to note that simian HAV Macaca/Germany/2021 only caused increased liver enzyme activities in Mauritius-origin cynomolgus macaques. It is possible that Mauritius-origin monkeys are more sensitive to factors causing liver enzyme elevation, as Mauritius-origin cynomolgus macaques generally show a 20% to 40% higher liver enzyme activity when compared to Asia mainland-origin animals (*20*). It is also possible that simian HAV Macaca/Germany/2021 circulates in cynomolgus monkeys from Asia generating immunity of Asia-origin animals opposed to HAV-naïve Mauritius-origin macaques.

## CONCLUSION

We conclude that simian HAV Macaca/Germany/2021 is present in cynomolgus macaques and that it poses a potential risk particularly for Mauritius-origin captive monkeys that are intended for use in scientific purposes. In order to prevent infection and associated elevations in liver enzyme activity, it is recommended to vaccinate animals. A commercial vaccine against human HAV has demonstrated efficacy in this matter.

## ACKNOWLEDGEMENT

We acknowledge and highly appreciate the work of Dr. Artur Kaul and Dr. Annette Dammert from German Primate Center in Goettingen, Germany, for establishing and conducting RT-PCR assays for simian HAV Macaca/Germany/2021. We also acknowledge the contribution from David Dahlhaus who formerly worked for Labcorp and participated in the root cause analysis.

## About the author

Mr. Mecklenburg is a veterinary pathologist at Labcorp Early Development Services GmbH, Muenster, Germany. His primary research interests include toxicologic pathology, health risk assessment, and management of non-human primates under laboratory conditions.

