## Appendix for "A New Variant of Hepatitis A Virus Causing Transient Liver Enzyme Elevations in Mauritius-origin Laboratory-housed Cynomolgus macaques": Appendix.pdf

### Description of the sequence-independent single-primer amplification approach.

Viral RNA was extracted from blood samples using the QIAamp Viral RNA kit. We then treated 30 µL of each viral extract with 20 µL of Turbo DNase master mix and incubated at 37°C for 30 minutes. After cleaning and concentrating RNA with a Zymo clean-up kit, we added 1 µL of primer A stock to each 4 µL of extracted RNA, and then heated at 65°C for 5 minutes, followed by cooling at 4°C for 5 minutes. Next, we prepared a master mix of 2 µL 5X RT buffer, 1 µL 10 mM dNTP, 1 µL water, 0.5 µL 0.1M DTT, and 0.5 µL SSIV reverse transcriptase per sample, and added 5 µL of master mix to each cleaned extract. We then ran the reverse transcription at 42°C for 10 minutes. After the 10 minute incubation, we added 5 µL consisting of 1 µL 5X Sequenase buffer, 3.8 µL water, and 0.15 µL Sequenase to each sample, and incubated at 37°C for 8 min. After the 8 minute incubation, we added 0.6µL consisting of 0.45 µL Sequenase dilution buffer, and 0.15 µL Sequenase and incubated at 37°C for an additional 8 minutes. Each cDNA library was then amplified using a master mix of 5 µL AccuTaq LA 10x Buffer, 2.5 µL dNTP mix, 1µL DMSO, 0.5 µL AccuTaq LA DNA Polymerase, 35 µL nuclease free water, and 1 µL Primer B. Each reaction ran on a thermocycler with 98°C for 30 seconds, followed by 30 cycles of 94°C for 15 seconds, 50°C for 20 seconds, and 68°C for 2 minutes, a final step of 68°C for 10 minutes, and then a 4°C hold. Finally, each amplified cDNA library was cleaned with a 1:1 ratio of AMPure XP beads. Beads were added and incubated at room temperature for 10 minutes, after which point the libraries were placed on a magnet to hold sequences intact while the supernatent was removed. Beads were then washed twice with 70% ethanol, where ethanol was removed after each wash, air dried, and then resuspended in 48 µL of water. After this DNA library preparation, all samples were sequenced on an Illumina NovaSeq6000 instrument at the University of Wisconsin BioTechnology Center.

**Primer pairs used in qRT-PCR for detection of simian HAV Macaca/Germany/ 2021.**

| name | pool | seq | size | %gc | tm (use 65) |
|---|---|---|---|---|---|
| **6488-HAV_1_LEFT** | 1 | TGGTGAGGGGACTTGATACCTC | 22 | 54.55 | 61.08 |
| **6488-HAV_1_RIGHT** | 1 | TGTGGATAAACGGTAAGGGAAGC | 23 | 47.83 | 60.87 |
| **6488-HAV_2_LEFT** | 2 | ACAATGAGCAATTTGCAGTGCA | 22 | 40.91 | 60.40 |
| **6488-HAV_2_RIGHT** | 2 | GATACCAACCTGAGGGTCTGGA | 22 | 54.55 | 61.08 |
| **6488-HAV_3_LEFT** | 1 | TCCAATGTTGCATCTCATGTCAGA | 24 | 41.67 | 60.89 |
| **6488-HAV_3_RIGHT** | 1 | TCTAGTGTCTGCCGAAAGACCT | 22 | 50.00 | 61.00 |
| **6488-HAV_4_LEFT** | 2 | AAATGAAATTCTGCCCCCTCCC | 22 | 50.00 | 61.08 |
| **6488-HAV_4_RIGHT** | 2 | TTTGATGAACCCGAGAGATGCA | 22 | 45.45 | 60.21 |
| **6488-HAV_5_LEFT** | 1 | AGAGGCTGACCAATTCTGTGTG | 22 | 50.00 | 60.73 |
| **6488-HAV_5_RIGHT** | 1 | ACTCCCAAGGCATTCATAACCC | 22 | 50.00 | 60.54 |
| **6488-HAV_6_LEFT** | 2 | AGATGCTGATCCTGCTGAGTCT | 22 | 50.00 | 61.14 |
| **6488-HAV_6_RIGHT** | 2 | TGGACCTCAATTCATCTTTAGGACA | 25 | 40.00 | 60.14 |
| **6488-HAV_7_LEFT** | 1 | GAGCTCCAGGTATTGATGCCAT | 22 | 50.00 | 60.41 |
| **6488-HAV_7_RIGHT** | 1 | GAAAATTTGTTTAAGCAAGTCATGAAAGG | 29 | 31.03 | 60.02 |

**Histology of liver from a colony animal with increased ALT and GLDH enzyme activity. H&E stain. Minimal mononuclear cell infiltrates in peri-portal region with very few degenerating hepatocytes (200x).**

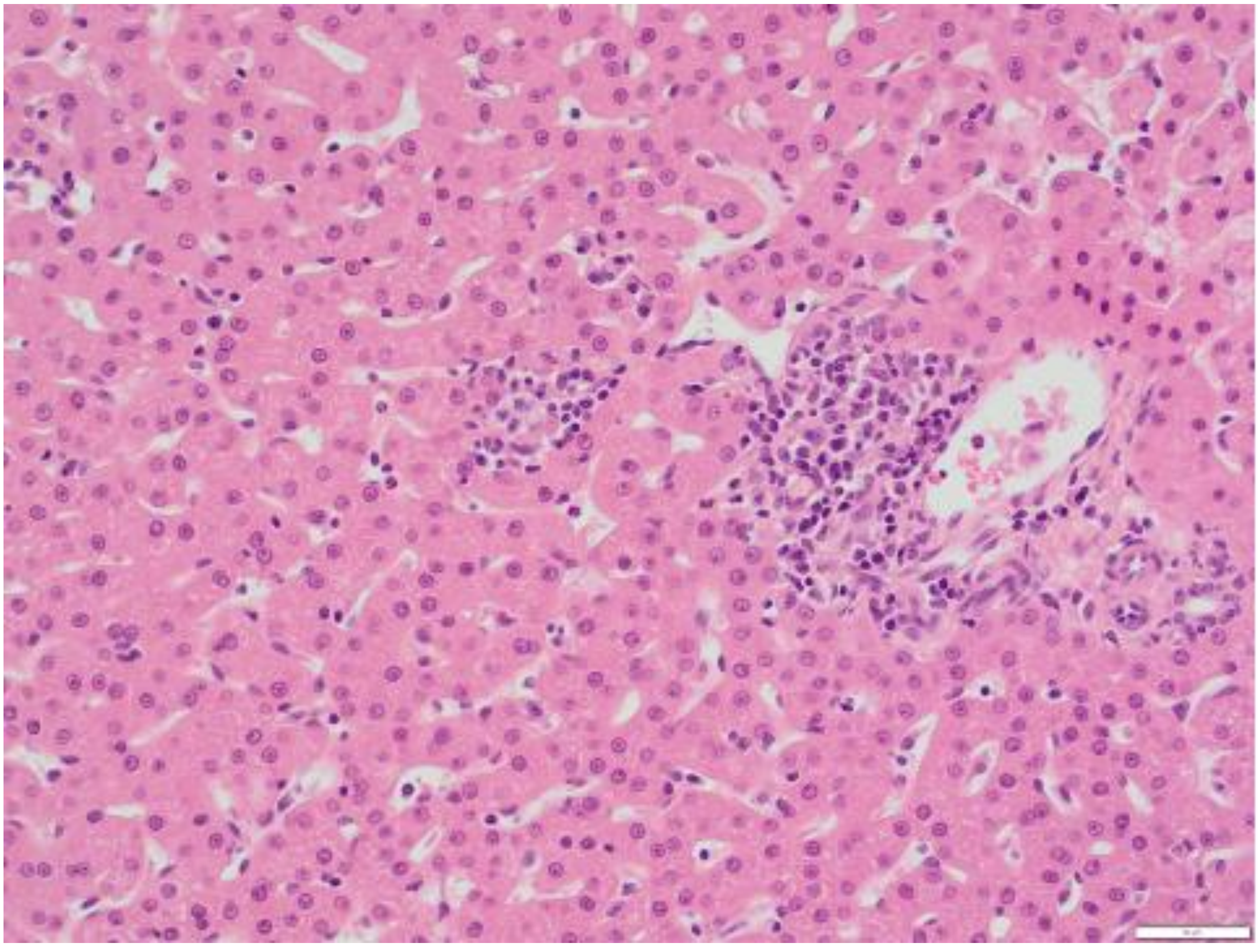

**Histology of liver from a colony animal with increased ALT and GLDH enzyme activity. H&E stain. Single cell necrosis of hepatocytes (400x).**

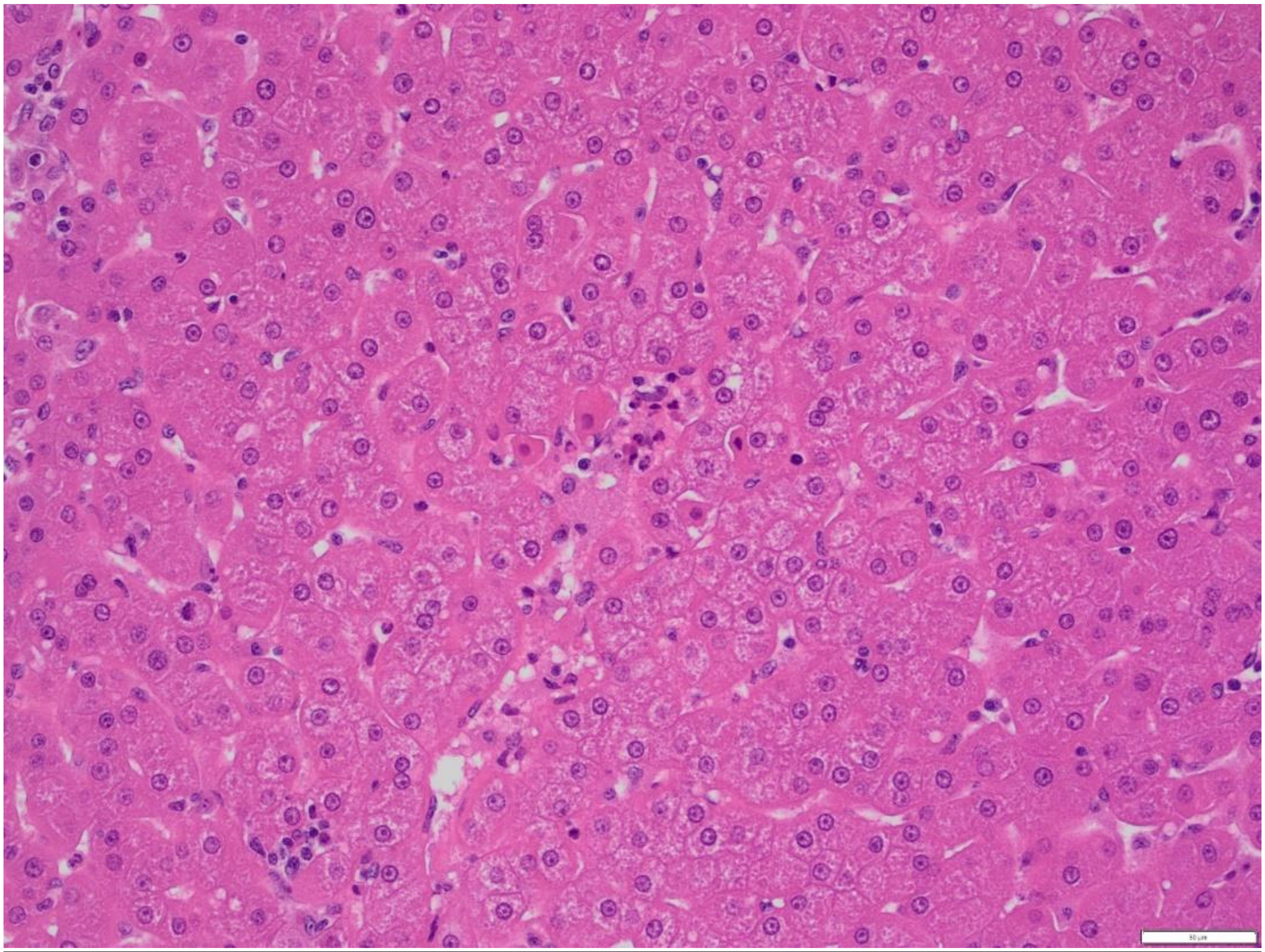
